# Comparative analysis of public RNA-sequencing data from human intestinal enteroid (HIEs) infected with enteric RNA viruses identifies universal and virus-specific epithelial responses

**DOI:** 10.1101/2021.03.30.437726

**Authors:** RJ Cieza, JL Golob, JA Colacino, CE Wobus

## Abstract

Acute gastroenteritis (AGE) has a significant disease burden on society. Noroviruses, rotaviruses and astroviruses are important viral causes of AGE but are relatively understudied enteric pathogens. Recent developments in novel biomimetic human models of enteric disease are opening new possibilities for studying human-specific hostmicrobe interactions. Human intestinal enteroids (HIE), which are epithelium-only intestinal organoids derived from stem cells isolated from human intestinal biopsy tissues, have been successfully used to culture representative norovirus, rotavirus and astrovirus strains. Previous studies investigated host-virus interactions at the intestinal epithelial interface by individually profiling the epithelial transcriptional response to a member of each virus family by RNA sequencing (RNA-seq). We used these publicly available datasets to uniformly analyze these data and identify shared and unique transcriptional changes in the human intestinal epithelium upon human enteric virus infections.

## Introduction

Acute gastroenteritis (AGE) is a major source of illness globally. It is defined as the inflammation of the stomach and intestines accompanied by rapid onset diarrhea, nausea, vomiting and abdominal pain. Five major virus families have been identified as etiological agents of viral gastroenteritis: noroviruses, sapoviruses (both viruses single-stranded positive-sense RNA viruses belonging to the *Caliciviridae* family), rotaviruses (double-stranded RNA viruses belonging to the *Reoviridae* family), adenoviruses (double-stranded DNA viruses belonging to the *Adenoviridae* family) and astroviruses (single-stranded RNA viruses belonging to the family *Astroviridae*) [1]. Only 20 – 30% of AGE cases in the United States have a specific causal virus identified [2]. Globally, noroviruses have the highest prevalence across all age groups, causing one fifth of all AGE cases [3]. Rotaviruses are the leading cause of AGE in children <5 years of age despite the availability of a vaccine [4]. Human astroviruses typically cause diarrhea in children <2 years of age but extraintestinal disease, including meningitis/encephalitis, is observed in immunocompromised individuals [5]. A full understanding of viral AGE intestinal pathogenesis has been challenging due to genetic diversity within each viral family and due to the lack of suitable experimental models resembling the complexity of the human intestinal epithelium.

Human intestinal enteroids (HIEs), are patient-derived endoderm-only 3D structures composed of heterogeneous cell populations recapitulating intestinal tissues in vivo, providing a more faithful experimental model than immortalized and transformed cells. A large degree of epithelial cellular diversity is observed in HIEs: Columnar intestinal epithelial, stem/progenitor, enteroendocrine and secretory cells can be observed depending on the culture conditions [6]. Spheroid structures can also be transitioned into 2D monolayers to facilitate host-microbe interaction studies in physiologic asymmetric oxygen conditions [7]. Furthermore, HIEs have been used to study the pathogenesis of several enteric viruses including difficult-to-cultivate ones [8, 9].

HIEs provide a uniquely complex system to study host-virus interactions at the intestinal epithelial interface and have been used to profile the epithelial transcriptional response to viral gastroenteritis by RNA sequencing (RNA-seq) in several studies. Non-classic astrovirus (VA1 strain) upon infection of duodenal-HIEs triggered type I and type III interferon (IFN) signaling [10]. Ileal-HIEs infected with human norovirus (GII.4 strain) showed high enrichment in STAT1 and STAT2 binding sites, suggesting JAK-STAT signaling pathway activation due to type I IFN signaling [11]. Jejunal-HIEs when infected with rotavirus (strain Ito) showed a predominant and conserved type III IFN response [12]. Integrating the conclusions from these studies that explore the pathogenesis of astrovirus [10], norovirus [11] and rotavirus [12] in HIEs is challenging since there are technical differences between studies. Each study uses HIEs grown from different tissue donors and varies the time-interval at which the host-response to the virus was evaluated. Furthermore, different library construction protocols can result in different sequence coverage, complexity and evenness [13]. Despite the limitations to comparative analysis of RNA-seq datasets generated from different studies, re-analysis of published public datasets can provide novel insights into a scientific question as long as the researchers take the biological context into consideration in which each dataset was generated [14].

Here, our goal was to re-analyze three publicly available datasets from HIEs infected with three representative viruses that cause AGE to gain further insights into whether there are shared and unique transcriptional changes in the intestinal epithelium upon virus infection from a qualitative assessment standpoint.

## Results

### Samples across datasets showed divergence due to tissue origin of the HIEs line and viral infection

The RNA-seq datasets from each one of the studies that we analyzed [10–12] were all generated from HIEs grown from biopsy tissue from the small intestine. However, in each study a different section of the small intestine from a different donor was used, specifically the terminal ileum (TI006 and TI365 lines) for norovirus infection [11], jejunum (J2 and J11 lines) for rotavirus infection [12] and duodenum (D124) for astrovirus infection [10] **(Supplemental Table 1)**. Hence it was not surprising that the main source of variance when looking at the three datasets together was not the treatment (mock-treated versus virus-infected) but rather the segment of the small intestine used to derive HIEs **(Figure 1A)**. In the datasets where more than one HIE line was used, the driver of variance still was the HIE line used in the experiment (although that is linked with the institution where the RNA-seq experiment was done). Still, when looking at each dataset individually and comparing between samples infected or not, consistent differences can be observed **(Figure 1B – D)**. Dimensional reduction via principal component analysis (PCA) showed similar results, where the primary source of variance when looking at all the datasets together was the tissue origin of the small intestine used to derive HIEs **(Supplemental Figure 1)**.

**Figure 1.**
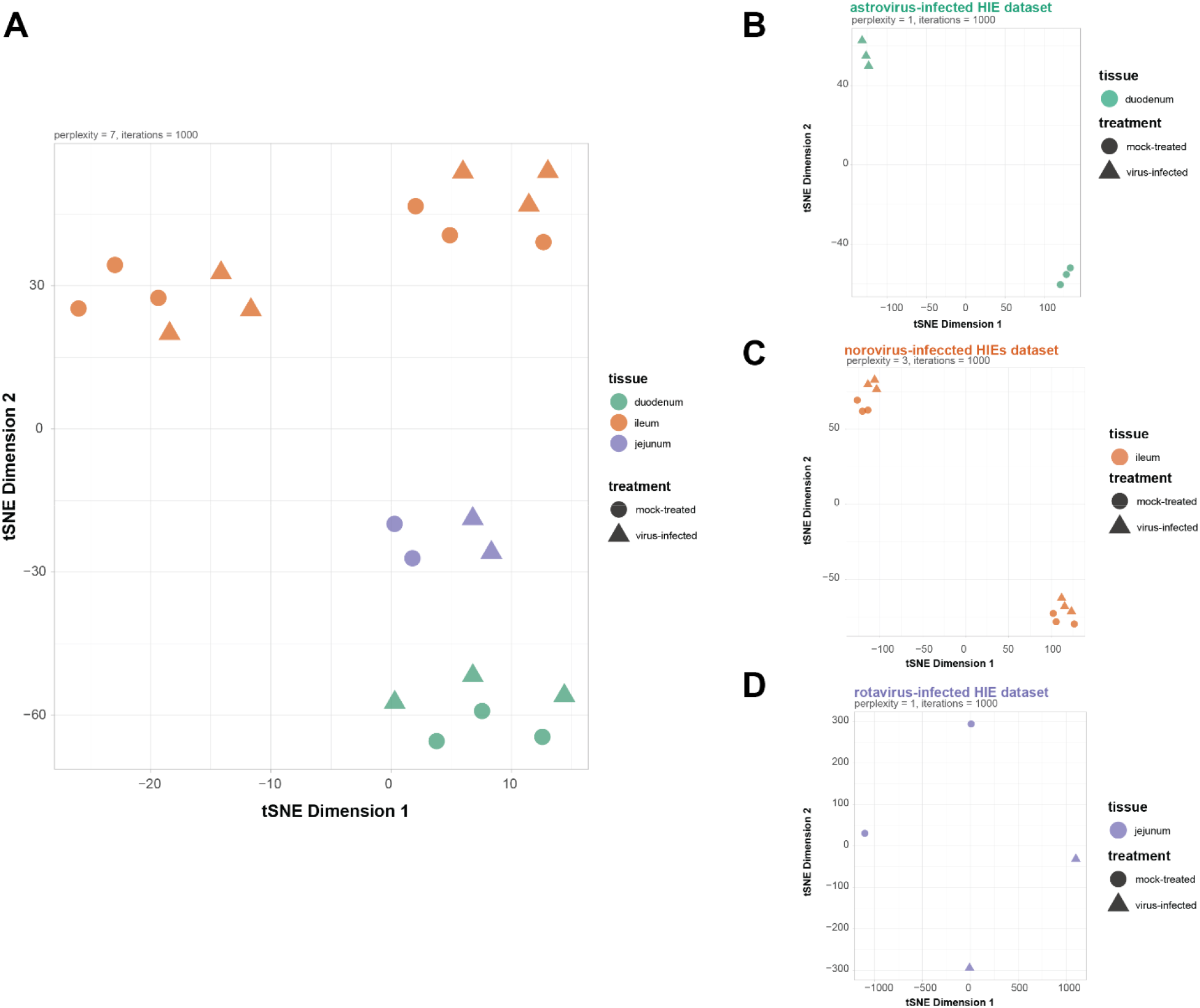
Samples across the three datasets showed divergence due to tissue origin of the HIE line and due to viral infection when looking at each dataset individually. Nonlinear dimensionality reduction with t-SNE of the 16 samples belonging to the three datasets analyzed. Samples are colored by the segment of the small intestine that was used to grow HIE and each one of the datasets analyzed correspond to a different small intestinal segment (duodenum, ileum and jejunum). The t-SNE plot shows that clustering is primarily driven by the tissue origin of HIE **(A)**. Nonlinear dimensionality reduction with t-SNE for each individual dataset (6, 12 and 4 samples for astrovirus-infected **(B)**, norovirus-infected **(C)** and rotavirus-infected **(D)** HIEs datasets respectively) shows clustering of the samples due to treatment (mock-treated versus virus-infected) as well as due to HIE line since in the norovirus-infected and rotavirus-infected HIE datasets more than one HIE line were used. Plots were generated with Rtsne and perplexity values were chosen for each plot after stability.

### Most non-weakly expressed genes are shared across datasets including a number of innate immune response genes

We next focused on finding differences and similarities in HIE transcriptional changes upon enteric viral infection between the three RNA-seq datasets analyzed. After filtering we considered 11824, 12917 and 12416 genes from astrovirus-infected duodenal-HIEs, norovirus-infected ileal-HIEs and rotavirus-infected jejunal-HIEs, respectively. The union for the three RNA-seq datasets consisted of 13581 genes with 82.92 % of these (11262 genes) present in all the datasets **(Figure 2A)**. Among the non-weakly expressed genes detected in all RNA-seq datasets were several IFN receptor genes (IFNLA, IFNGR and IFNLR) as well as IFN-stimulated genes. Other key players of the innate immune response such as genes associated with Toll receptor signaling (TLR1 – 3) **(Figure 2B)** and chemokines secreted in response to IFN such as CXCL10 and CXCL1 (data not shown) were also detected in all the datasets. Several of the selected shared genes annotated in the Entrez Molecular Sequence Database System as part of IFN responses showed estimated gene counts above 1000 (log_2_ values > 10), however, no genes coding for IFN were found in the list of genes identified in all RNA-seq datasets, except for IFN-λ (IFNL1-3) in rotavirus-infected jejunal-HIEs and IFN-ε (IFNE) which was partially shared across two of the RNA-seq datasets **(Supplemental Table 2)**. While biologically IFNs clearly play a critical role in limiting astrovirus [10] and norovirus infections [15], RNA sequencing could have limitations in sensitivity due to sequencing depth [16], which may explain this discrepancy. Altogether, when looking at the overall expression patterns across the different RNA-seq datasets a large number of the genes identified were shared across the three studies.

**Figure 2.**
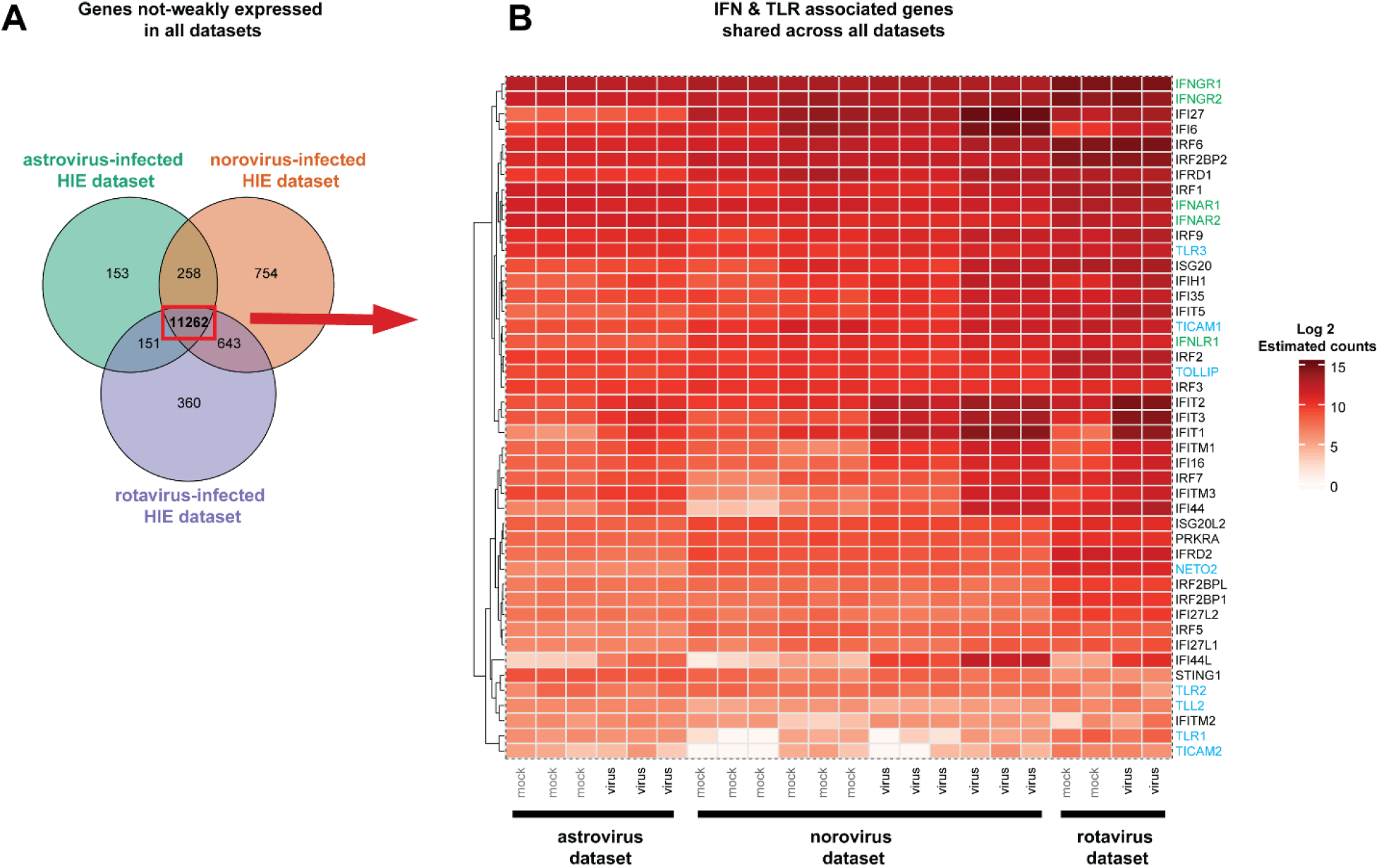
A large degree of overlap of non-weakly expressed genes is observed across all datasets. Estimated gene count matrices for each RNA-seq dataset were filtered to remove genes with low constant expression levels (non-weakly expressed) with the package HTSFilter in R. Log transformed estimated counts for selected genes (IFN and Toll receptor signaling associated) are shown. A total of 11262 genes were identified in the three RNA-seq datasets with norovirus-infected ileal-HIEs producing the largest number of genes identified in a single dataset **(A)**. Genes annotated in the Entrez Molecular Sequence Database System as part of IFN responses or involved in Toll receptor signaling detected in the three datasets are shown. Individual replicates (mock-treated samples in grey and virus-infected samples in black) for each study are shown. Additionally, highlighted in green are genes coding for IFN or IFN receptors and in blue are genes associated with Toll receptor signaling **(B)**

### Differential expression analysis showed that HIE responses against astrovirus, norovirus and rotavirus is primarily characterized by an up-regulation of interferon-stimulated genes (ISGs)

To determine the differentially expressed genes (DEGs) between infected versus mock controls in each RNA-seq dataset, we re-analyzed each dataset individually for differential expression with the package DeSeq2 to find whether the list of DEGs we obtained in this analysis was similar to what was previously published [10–12]. For each individual RNA-seq dataset, DEGs were identified among non-weakly expressed genes **(Table 1)** based on the criteria of an actual fold-change of at least 1.5 (log_2_-FC > 0.58) with a false discovery rate (FDR) cutoff of 1% (adjusted p-value < 0.01) using an Empirical Bayes (EB) approach with the R package ashr [17]. Relatively stringent FDR cutoffs were used to focus only on the strongest biological effects upon virus infection in each one of the datasets analyzed.

**Table 1:** Differential expression analysis summary. Differential analysis of RNA-seq data was performed on the three published RNA-seq datasets. Listed are the total number non-weakly expressed genes that were analyzed in each dataset to determine differentially expressed genes (DEG).

| **Table 1. Differential expression analysis summary** |  |  |  |  |  |
| --- | --- | --- | --- | --- | --- |
| **dataset** | **HIEs** | **# genes analyzed** | **DEGs** | **up-regulated-DEGs** | **down-regulated DEGs** |
| **astrovirus-infected** | **duodenal-HIEs** | **11824** | **42** | **42** | **0** |
| **norovirus-infected** | **ileal-HIEs** | **12917** | **109** | **109** | **0** |
| **rotavirus-Infected** | **Jejunal-HIEs** | **12416** | **75** | **71** | **4** |

The astrovirus-infected duodenal-HIEs revealed 42 DEGs out of a total of 11824 non-weakly expressed genes. The number of DEGs identified in this dataset was the smallest of the three RNA-seq datasets analyzed and all were up-regulated genes **(Figure 3A)**. When re-analyzing the norovirus-infected ileal-HIEs dataset, we evaluated the sequencing data from both HIE lines (TI006 and TI365) together since we wanted to explore universal patterns of host-response against norovirus GII.4. Additionally, sequencing data for both HIE lines were generated within the same institution for the same study and both HIE lines were grown from ileal biopsies. We found that of a total of 12917 non-weakly expressed genes, 109 were DEGs, which were all up-regulated **(Figure 3B)**. Finally, in the RNA-seq dataset generated from rotavirus-infected jejunal-HIEs, two jejunal-HIE lines were used to identify DEGs. In this study, out of a total of 12416 non-weakly expressed genes, 75 were DEGs with 71 up-regulated and 4 down-regulated genes **(Figure 3C)**.

**Figure 3.**
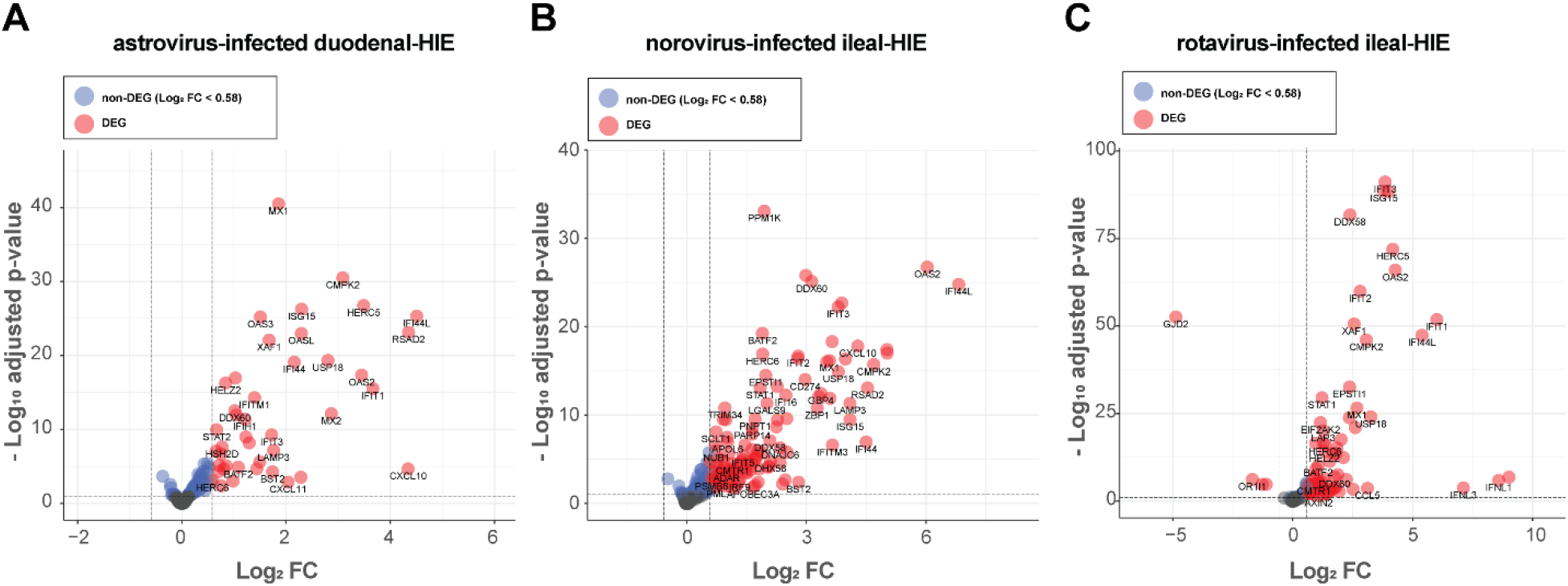
Differential expression analysis in HIEs infected with astrovirus, norovirus and rotavirus. Differentially expressed genes (DEGs) for all the three RNA-seq datasets re-analyzed are shown. DEGs are shown red (Log2 foldchange (FC) > 0.58 when compared to mock-treated samples and an FDR of less than 1%). In blue are shown genes with an FDR of less than 1% (adjusted p-value < 0.01) that were below the log2 FC threshold. FDR was calculated using an Empirical Bayes (EB) approach with the R package ashr. DEGs were identified in astrovirus-infected duodenal-HIEs **(A)**, norovirus-infected ileal-HIEs **(B)** and rotavirus-infected jejunal-HIEs **(C)** using the package DESeq2 and plotted using the package EnhanceVolcano in R.

A large degree of similarity was found in the list of DEGs identified in our analysis with what was previously published for each study [10–12]. The differences in the number of DEGs detected between our and published analysis are likely due to the different filtering methods applied to the matrices containing the estimated gene counts and the different log_2_ FC cutoff applied.

### Antiviral defense and IFN signaling represent a conserved response of the epithelium to viral infection

To identify shared DEGs, we compared the list of DEGs identified for each individual RNA-seq dataset. There were 33 DEGs common to all three datasets, suggesting they are part of a shared response against viruses by the intestinal epithelium **(Figure 4A)**. 25 out of the 33 common DEGs are annotated as part of the IFN response in the Entrez Molecular Sequence Database system **(Figure 4B)**. Of these, a number of IFN-stimulated genes (ISGs) were identified as part of the universal HIEs response to the three viruses, including IFN-induced proteins (IFIT1, IFIT2 and IFIT3) as well as signal transducer and activator of transcription protein 1 and 2 (STAT1 and STAT2). Shared DEGs were also evaluated in the Interferome database showing that 16 out of the 33 DEGs shared across RNA-seq datasets were associated with type I, II, III IFN responses **(Table 2)**. However, within the shared DEGs, virus-specific induction patterns were also observed. For example, we observed that the two ISGs 2’-5’-oligoadenylate synthetase (OASL), Interferon Induced Protein 44 (IFI44) and the Solute Carrier Family 15 Member 3 (SLC15A3), which potentiates MAVS- and STING-mediated IFN production [18], showed a stronger up-regulation in the norovirus-infected ileal-HIEs compared to astrovirus and rotavirus infection **(Figure 4B)**. On the other hand, IFN Induced Protein with Tetratricopeptide Repeats 1 (IFIT1) showed greater up-regulation in the rotavirus dataset versus either norovirus or astrovirus datasets. In addition, 30 genes were differentially expressed in only two out of the three datasets **(Figure 4C)**. These partially shared DEGs include for example C-X-C Motif Chemokine Ligand 11 (CXCL11) and Bone Marrow Stromal Cell Antigen 2 (BST2, also called Tetherin), which was up-regulated in response to norovirus and astrovirus but not rotavirus infection. A different pattern was observed for example for the transcription factors IFN Regulated Factor 7 and 9 (IRF7, IRF9), which were differentially expressed in the astrovirus and rotavirus but not norovirus datasets and the rotavirus and norovirus but not astrovirus datasets, respectively.

**Figure 4.**
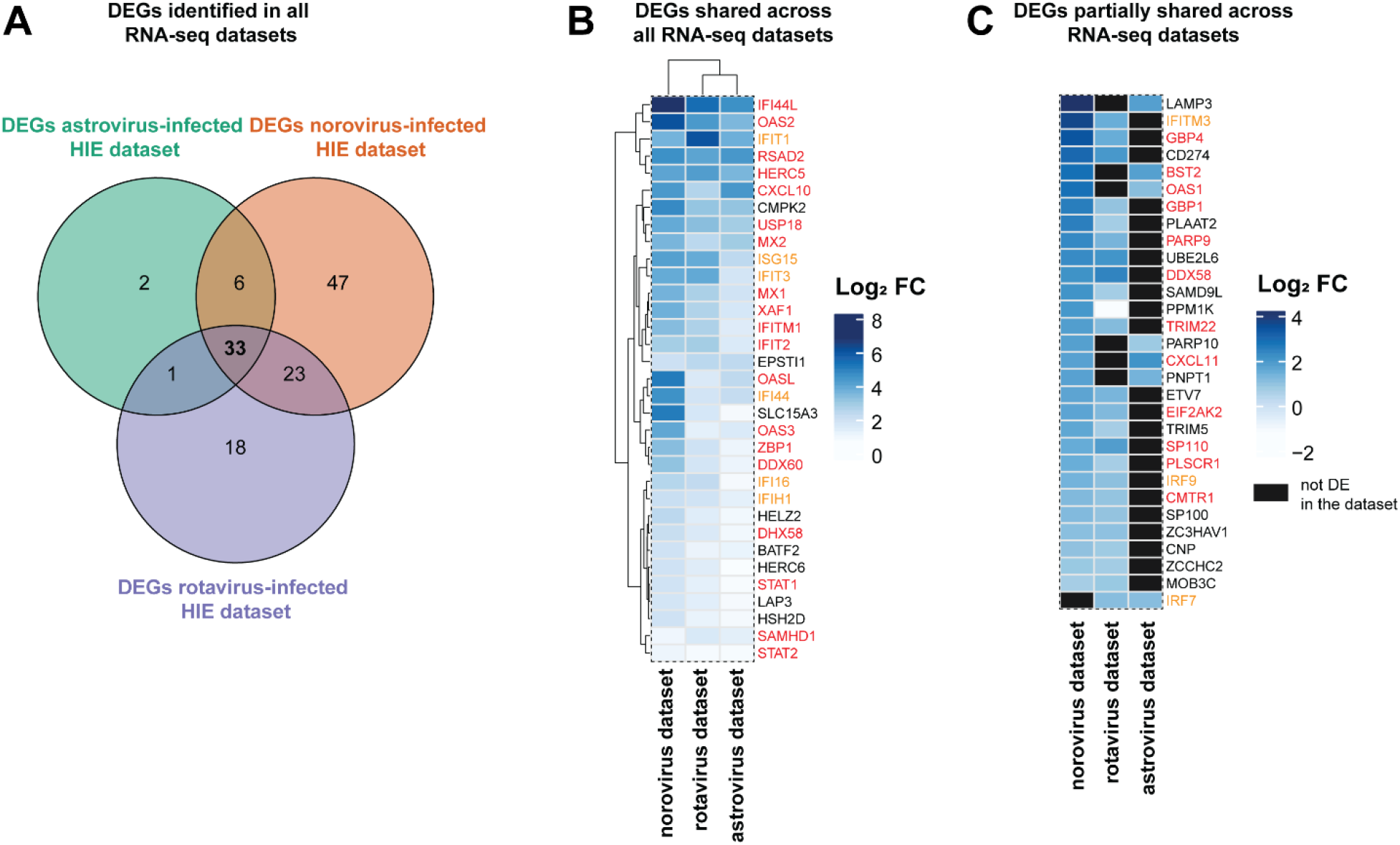
Shared differentially expressed genes (DEGs) across datasets, which are part of the conserved epithelial response to viral infection, are enriched in IFN associated genes. Shared and unique DEGs across all datasets were determined using the package DESeq2 and heatmaps generated with the package ComplexHeatmap in R. For a gene to be classified as differentially expressed it had to meet the criteria of an actual fold-change of at least 1.5 (log_2_-FC > 0.58) with a false discovery rate (FDR) cutoff of 1% (adjusted p-value < 0.01) using an Empirical Bayes (EB) approach with the R package ashr. The intersection and union of DEGs across the three RNA-seq datasets evaluated is shown **(A)**. Expression pattern for DEGs shared across all datasets is shown, with genes annotated in the Entrez Molecular Sequence Database System as part of IFN responses highlighted in red. Furthermore, in orange are highlighted genes with a symbol containing IFI (IFN inducible) or ISG (IFN-stimulated genes) **(B)**. Partially shared DEGs in two out of three RNA-seq datasets are also shown **(C)**. It can be seen that in the set of shared DEGs across the three datasets, there is an enrichment of genes associated with IFN responses.

**Table 2:**
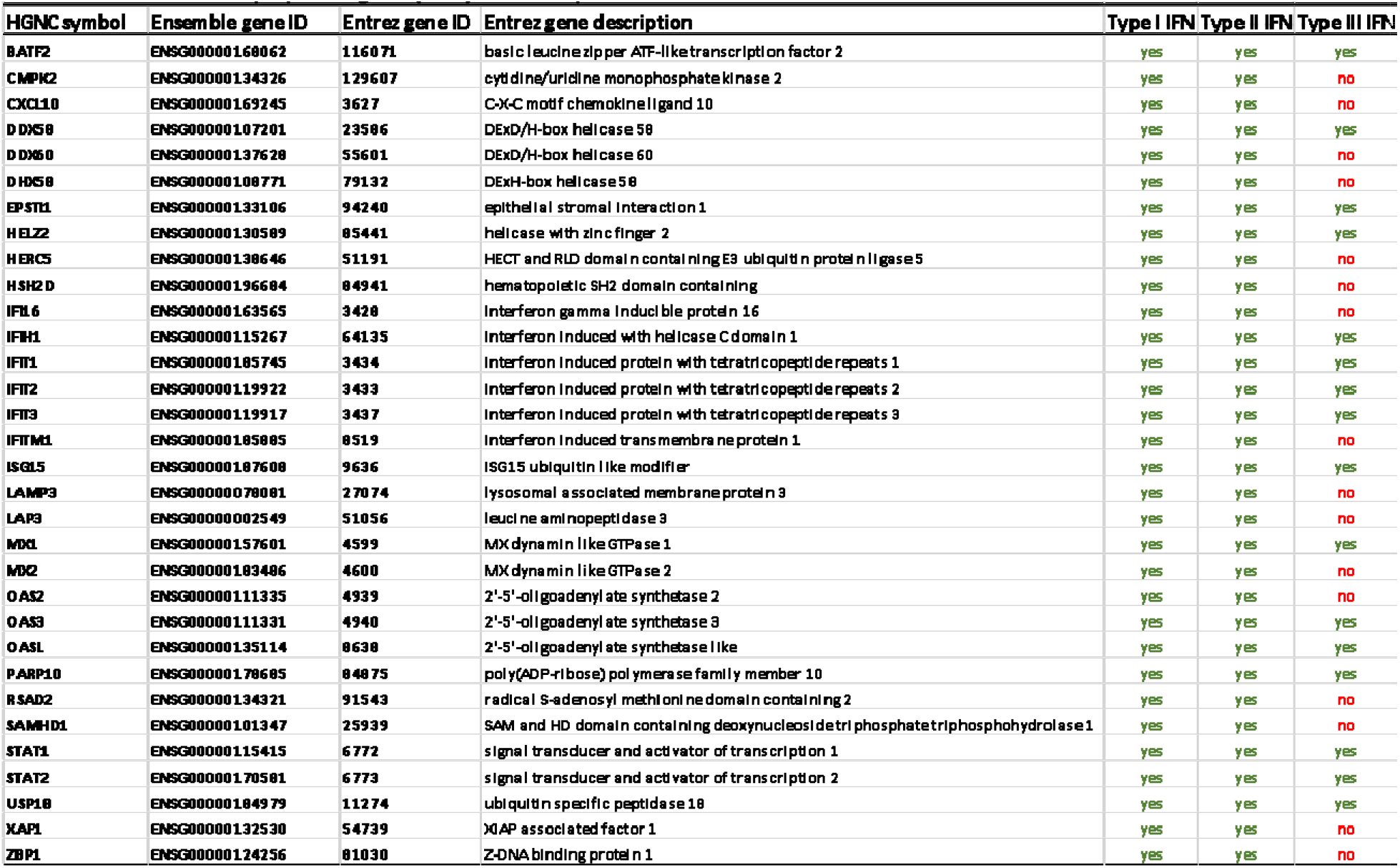
Differentially expressed genes (DEGs) shared in all datasets and IFN responses. DEGs that were identified in each dataset and likely part of a conserved response to astrovirus, norovirus and rotavirus in human intestinal enteroids (HIEs) as well as their association to Type I, II and III IFN responses.

In addition to specific genes, we also wanted to explore which biological processes were over-represented in the list of DEGs identified in each RNA-seq dataset. Towards that end we performed a gene ontology (GO) overrepresentation test comparing biological processes (BP). The dot plots highlight the top ten over-represented biological process (BP) categories identified in the list of DEGs for each one of the datasets **(Figure 5A - C)**. 18, 26 and 23 over-represented BP categories were found in astrovirus-infected, norovirus-infected and rotavirus-infected HIEs, respectively using an FDR cutoff of 0.1% (adjusted p value < 0.001), with the Benjamini–Hochberg (BH) step-up procedure **(Table 3)**. Ten over-represented BP categories were shared across all RNA-seq datasets **(Figure 5D)**. The top two over-represented BP categories were the same in the three datasets (“response to virus” [GO:0009615] and “defense response to virus” [GO:0051607]) with a gene ratio between 0.4 and 0.6, showing that close to half of the DEG identified for each dataset fall within both of these categories, independently of the infecting virus. In the next group of over-represented BP categories, we found several processes associated with a type I IFN response (“regulation of type I interferon production” [GO:0032479] and “type I interferon production” [GO:0032606]) with gene ratios close to 0.2. The total number of over-represented BP categories was similar across the three RNA-seq datasets, even though the number of DEGs identified in the astrovirus-infected duodenal-HIEs was much lower than in the other two datasets **(Table 3)**.

**Figure 5.**
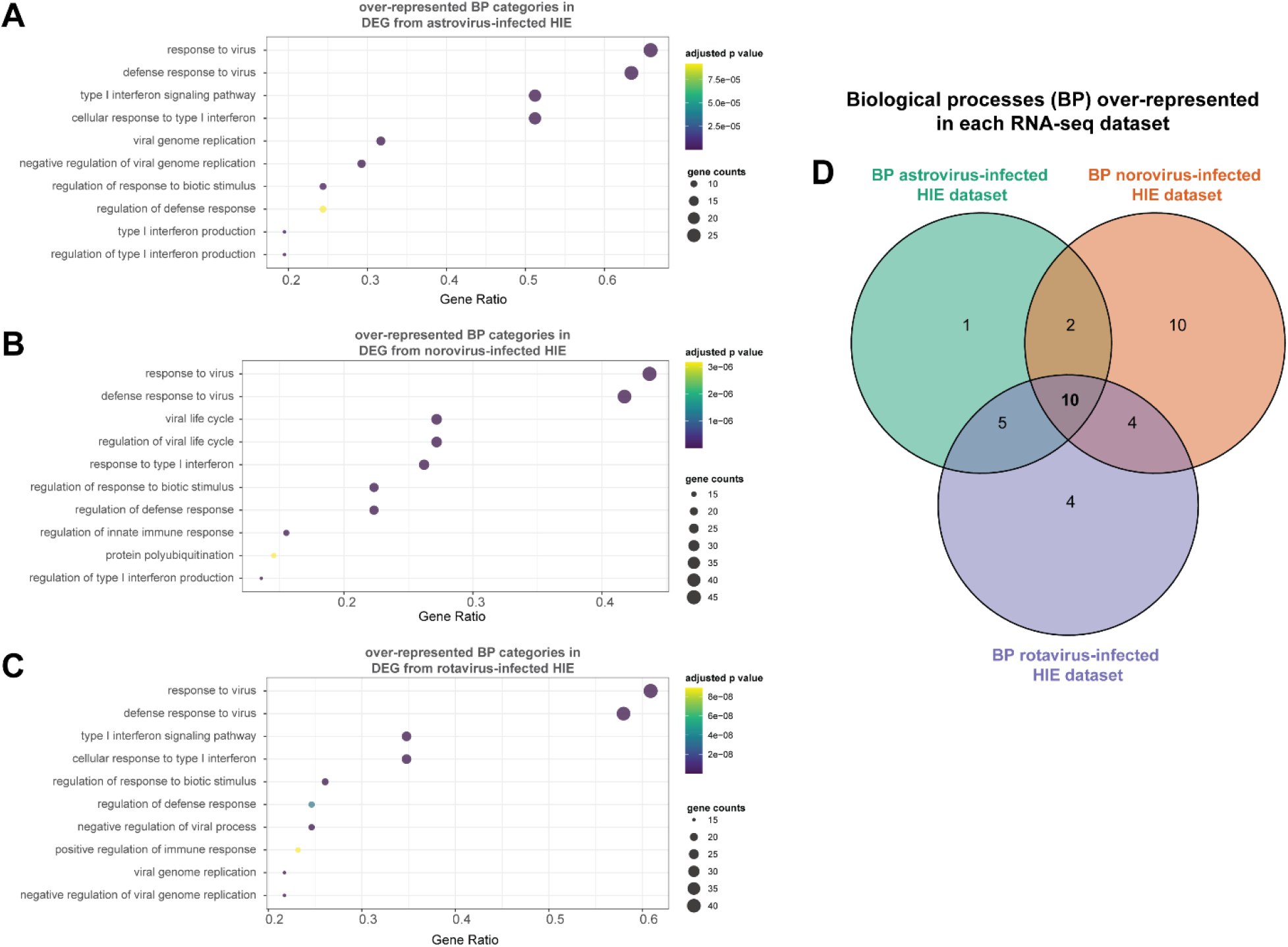
Gene Ontology (GO) over-representation analysis show that the top over-represented biological processes (BP) for all the datasets were associated with defense response to virus and type I IFN response. Gene ontology (GO) analysis was performed in the list of DEG identified for each one of the datasets analyzed independently to identify over-represented biological processes (BP). The top 10 over-represented BP categories are shown for each one the dataset as well as the number of genes contributing to the over-representation of each one of these BP. Astrovirus-infected duodenal-HIE **(A)**, norovirus-infected ileal-HIE **(B)** and rotavirus-infected jejunal-HIE **(C)** are shown. An adjusted p-value cutoff of 0.001 was used to determine significant over-represented BP. Shared and unique over-represented BP categories across all datasets were determined from the list of overrepresented BP categories obtained for each dataset **(D)**. Over-represented BP categories were identified with the package clusterprofiler in R.

**Table 3:**
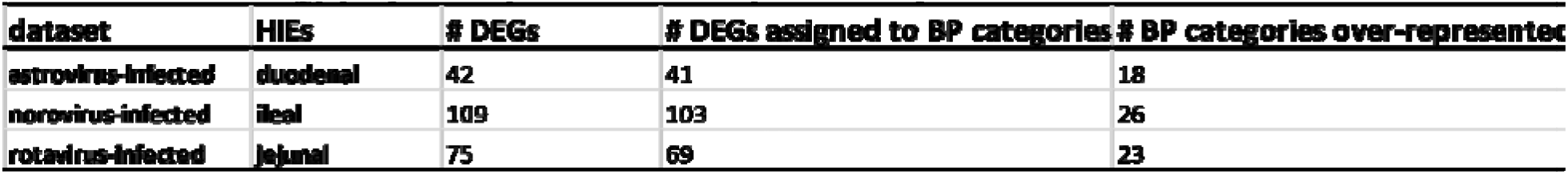
Gene Ontology (GO) over-representation analysis summary. Over-representation analysis of biological processes (BP) was performed with the package ClusterProfiler in R on three published datasets of virus-infected HIEs. An FDR cutoff of 0.1% (adjusted p value < 0.001), with the Benjamini–Hochberg (BH) step-up procedure was used to determine significant over-represented BP. Listed are the total number of DEGs identified for each dataset as well as the number of DEGs that were assigned to a BP after over-representation analysis. For each dataset the number of BP over-represented are also shown.

| **Table 3. Gene Ontology (GO) over-representation analysis summary** |  |  |  |  |
| --- | --- | --- | --- | --- |
| **dataset** | **HIEs** | **# DEGs** | **# DEGs assigned to BP categories** | **# BP categories over-represented** |
| **astrovirus-infected** | **duodenal** | **42** | **41** | **18** |
| **norovirus-infected** | **ileal** | **109** | **103** | **26** |
| **rotavirus-infected** | **jejunal** | **75** | **69** | **23** |

Overall, a shared set of DEGs informing a shared set of biological processes (BP) were identified in the three RNA-seq datasets. This conserved, predominant response of the intestinal epithelium to viral infection is mediated by antiviral defense and IFN signaling pathways.

### Distinct DEGs and biological processes (BP) were also noted for each enteric virus

In addition to shared or partially shared DEGs, we also identified 2, 47 and 18 DEGs uniquely identified in astrovirus-infected, norovirus-infected and rotavirus-infected HIEs, respectively **(Figure 4A, Supplemental Table 3)**. In the dataset generated from astrovirus-infected duodenal-HIEs, two DEGs (IFI27 and REV3L) not overlapping with the other datasets were identified. Interferon Alpha Inducible Protein 27 (IFI27) expression sensitizes cells to apoptotic stimuli [19], which may help the intestinal epithelium to repel astrovirus-infected cells. Dysregulation of REV3L the catalytic subunit of DNA polymerase ζ engages the innate immune response, with potential antiviral consequences [20]. However, additional studies are needed in the future to test these hypotheses. 47 DEGs were found exclusively in norovirus-infected ileal-HIE, including IFN-induced and -stimulated proteins IFI35 and IFIT5 as well as the innate immune receptor toll-like receptor (TLR) 3 and several tripartite motif (TRIM) proteins (TRIM 14, 21, 25, 34 and 56). Among the 18 DEGs found exclusively in the rotavirus-infected jejunal-HIE dataset were IFN-λ (IFNL1, IFNL2 and IFNL3).

In line with unique DEGs, there were 1, 4, and 10 over-represented BP that were uniquely identified in each one of the three datasets from astrovirus, rotavirus and norovirus datasets, respectively **(Figure 5D)**. However, the majority of these BP categories showed low gene ratios (gene ratio < 0.1). In astrovirus-infected duodenal-HIEs “regulation of defense response to virus” [GO:0050688] was the only uniquely over-represented BP category for the dataset **(Table 4)**. However, it was driven by genes not uniquely differentially expressed in this dataset (IFIT1, HERC5, DDX60, DHX58 and STAT1) **(Figure 6)**. A likely explanation is that the combination of these genes might play a more relevant part of the host response triggered in the intestinal epithelium against astrovirus than norovirus or rotavirus. The chemokine CCL5 and IFN-λ (IFNL1-3) were important for 3 out of the 4 BP overrepresented uniquely in rotavirus-infected HIEs **(Table 4)**. These genes were exclusively differentially expressed as part of the HIEs response to rotavirus **(Figure 6A)**. Norovirus infection upregulated the most unique BP categories with 8 out of 10 over-represented BPs including TRIM family proteins **(Table 4)**.

**Figure 6.**
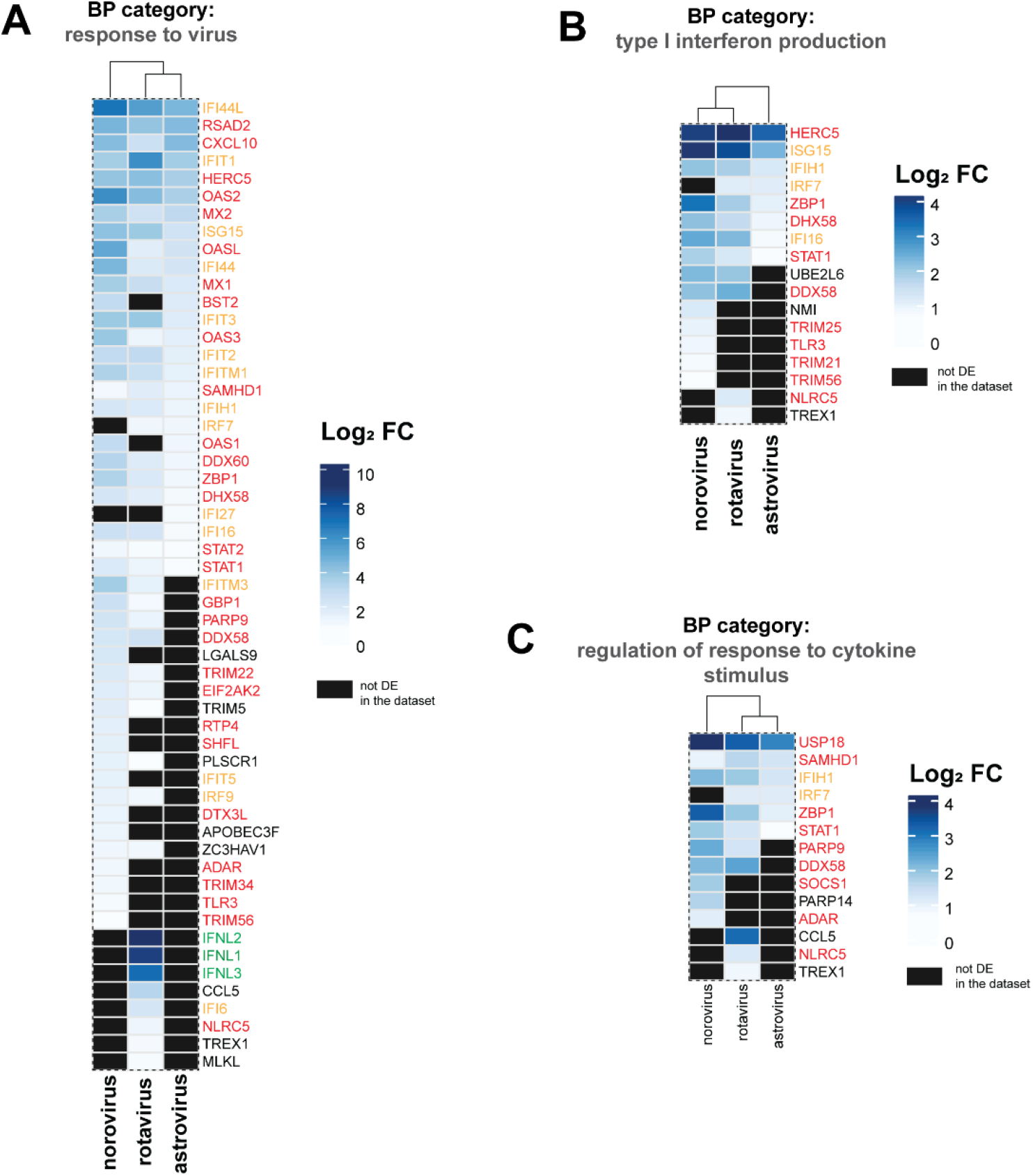
Expression patterns in shared over-represented biological processes (BP) in the three RNA-seq datasets after gene ontology (GO) over-representation analysis. Heatmap-like functional classification displays which differentially expressed genes (DEGs) were assigned to the BP categories “response to virus” **(A)**, “type I interferon production” **(B)** and “regulation of response to cytokine stimulus” **(C)**. For each dataset, in dark grey are genes that were not differentially expressed for the indicated RNA-seq dataset. Genes annotated in the Entrez Molecular Sequence Database System as part of IFN responses highlighted in red. Furthermore, in orange are highlighted genes with a symbol containing IFI (IFN inducible) or ISG (IFN-stimulated genes) and in green genes coding for IFN proteins. Log2 fold-change (FC) values, within the range of each dataset are shown. Over-represented BP categories were identified with the package clusterprofiler and heatmaps generated with the package ComplexHeatmap in R.

**Table 4:**
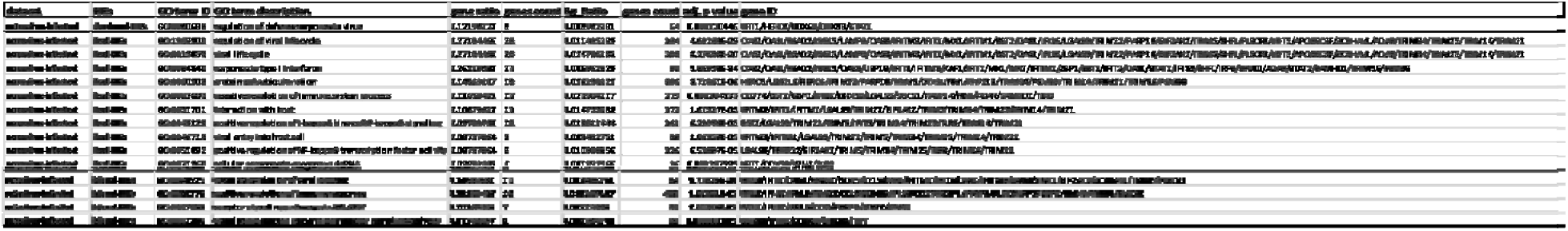
Over-represented BPs uniquely identified in each RNA-seq dataset. Over-representation analysis of biological processes (BP) with the package ClusterProfiler in R was used to identify uniquely over-represented BPs for each dataset. An FDR cutoff of 0.1% (adjusted p value < 0.001), with the Benjamini–Hochberg (BH) step-up procedure was used to determine significant over-represented BP categories.

While the top shared over-represented BP categories were the same in the three RNA-seq datasets **(Figure 5)**, we noted virus-specific differences in the expression pattern of DEGs assigned to each category **(Figure 6)**. For example, of the 55 genes included in the “response to virus” BP category, 23 were expressed in all datasets, while 13 were differentially expressed in two datasets, and the remaining 19 genes were only differentially expressed in one dataset **(Figure 6A)**. A similar pattern of shared, partially shared, and unique DEGs can be seen in other overrepresented BPs, for example type I IFN production **(Figure 6B)** and regulation of response to cytokine stimulus **(Figure 6C)**. In summary, shared and unique DEGs inform similar biological processes that are triggered in HIEs as part of an intestinal epithelial response to viral infection.

## Discussion

In this study, we wanted to identify universal and unique patterns of the intestinal epithelial response to enteric viral infection to contribute to a better understanding of the pathophysiology of acute gastroenteritis (AGE). Our meta-analysis draws from studies done at different centers, using HIEs from both different regions in the gut and distinct donors. This biological variability emphasizes the robustness of our ultimate finding, which was that IFNs and ISGs are central to the innate immune response to viral infections within the gut.

Our analysis was aware of the caveats presented from combining distinct RNA-seq datasets (a) sequenced at different institutions (b) obtained from HIEs grown from different sections of the small intestine (jejunum, duodenum and ileum), and (c) collected at different times post-virus infection (6, 24 and 48-hours post-infection with rotavirus, astrovirus and norovirus, respectively). These differences were noticeable when the data was subjected to dimensionality reduction, where t-Distributed Stochastic Neighbor Embedding (t-SNE) showed that the primary factor driving the differences between the three RNA-seq datasets was likely the anatomical origin of the biopsy used to generate HIEs **(Figure 1)**. Despite the variance across datasets due to the biopsy source used to grow HIEs, 89.92% of non-weakly expressed genes were identified in the three datasets (11262 out of 13581 genes). This was particularly useful for downstream differential analysis for each one of the RNA-seq datasets, since DEGs were identified from mostly similar lists of genes.

Several over-represented biological processes were part of type I IFN production, signaling and responses **(Figure 5)** and thus are part of the critical defense response in the intestinal epithelium to enteric viruses. This multigene family encodes 13 partially homologous IFNα subtypes, IFNβ and several other IFN gene products that are less well characterized [19]. However, in this study we were not able to detect either IFNα or IFNβ in any of the RNA-seq datasets. Only IFNε (IFNE) was detectable in the datasets generated from astrovirus-infected duodenal-HIEs and norovirus-infected ileal-HIEs, but IFNE was not differentially expressed. This is in contrast to our previously published research, where we found that colonic and ileal HIEs showed detectable IFNβ gene expression in response to astrovirus infection by quantitative PCR [10]. This highlights a caveat of transcriptomic analysis, whereby the findings are influenced by the depth of sequencing required to pick up, in this case, these specific IFN genes. For example, mammalian reovirus infected T84 intestinal cells and enterovirus 71-infected HT29 cells express lower levels of IFNβ transcripts compared to IFNλ [21, 22]. As such the level of IFNα and/or IFNβ production in the HIEs could have been below the limit of detection of RNA-sequencing. Similarly, we observed that a type III IFN response was a significant part of the epithelial response to rotavirus, with IFNλ (IFNL1, IFNL2 and IFNL3) being differentially expressed, consistent to what the authors originally reported [12]. Changes in IFNλ however were not subjected to differential expression analysis, since these genes had very low relative expression levels and were thus filtered out by our stringent analysis criteria. Thus, a greater sequencing depth would have been required to pick up these IFNλ genes since our research on astroviruses [10] was able to detect IFNλ by quantitative PCR in colonic and ileal-HIEs at 24 hours post-infection. Nevertheless, it is also possible that the production of IFNλ by HIEs in response to rotavirus might be of a larger magnitude than in response to norovirus or astrovirus infections. Comparative infections with all viruses in the same HIE line would be required to test this possibility in the future.

Type I and III IFN production and signaling typically occurs in response to stimulation of pattern recognition receptors (PRRs), including toll-like receptors (TLRs). We found TLR3 to be the only TLR differentially expressed and only in ileal HIEs in response to norovirus (≈ fold-change of 2). While TLR1, 2 and 3 were detectable across all datasets, they were only differentially expressed in response to norovirus infection. TLR3 is primarily present in the endosomal compartment and senses double-stranded (ds) RNA [22, 23]. DsRNA is a RNA virus replication intermediate present during infection with all three viruses. Thus, TLR3 could conceivably function in the antiviral response to all three viruses. However, TLR3 may only be differentially expressed in the norovirus-infected HIEs dataset due to the timing of sample collection (48 hours post-infection) since an amplification of the baseline expression level of TLR3 during viral infection of intestinal epithelial cells is part of the ISG response [22]. It is also conceivable that TLR3 might play a more unique role in the epithelial response to human norovirus rather than astrovirus and rotavirus. There is precedent from murine norovirus that TLR3 contributes to controlling murine norovirus infection in vivo [24]. In contrast, a number of ISGs, including IFN-stimulated gene 15 (ISG15), a central player in the host antiviral response [25], and the E3 ubiquitin-protein ligase HERC5, were strongly up-regulated in all datasets and classified as part of the type I IFN production biological process.

Overall, while comparative analysis of dissimilar RNA-seq datasets from the literature has limitations, our processing pipeline and filters (removal of weakly-expressed genes, higher FDR cutoffs) limited these caveats and allowed us to draw several conclusions. Specifically, we identified key shared players across all datasets, which are part of the response to virus (IF44, IF44L, STAT1, STAT2, MX1 and MX2), demonstrating that HIEs employ a generic antiviral response. However, this is augmented by more tailored responses depending on the virus (e.g., IFI27 in response to astrovirus, and TLR3 in response to norovirus). We hope this comparative study provides a useful resource to the scientific community that stimulates further explorations into common and unique aspects of the host response to virus infections at the human intestinal epithelium. These studies should be complemented by comparative studies between different viral pathogens that cause gastroenteritis in the same host genetic background.

## Methods

### Cell lines and virus infection

Each public RNA-seq dataset was generated in a different institution with variations in the experimental protocol used for viral infection, which we briefly summarize here. To generate the RNA-seq dataset from duodenal-HIEs infected with astrovirus, HIEs were seeded in 48-well plates as 2D monolayers in complete L-WRN medium and infected with astrovirus strain VA1 at an MOI of 1 based on genome copies per cell for 1 hour at 37°C or mock. Total RNA was extracted for sequencing at 24 hours post-infection [10]. Ileal-HIEs monolayers grown in 48-well plates were incubated for 2 hours at 37°C with stool filtrates containing a patient-derived GII.4 human norovirus strain (~ 1 x 10^6^ viral RNA copies) or mock-treated. Samples were collected at 48 hours post-infection before sequencing [11]. For the RNA-seq dataset from rotavirus-infected jejunal-HIEs, spheroid (3D) HIEs were grown in complete media with growth factors (CMGF+) and differentiated for 3-4 days before infection. Then HIEs were infected with human rotavirus (HRV) strain Ito and sequenced at 6 hpi [12].

### Selection of GEO and EMBL-EBI data

The sequencing reads obtained from RNA-seq experiments in each one of the studies re-analyzed were obtained from the Gene Expression Omnibus (GEO) or the European Bioinformatics Institute (EMBL-EBI) ArrayExpress collection. For the norovirus-infected ileal-HIEs study FASTQ files from a total of 12 HIEs samples (6 mock-treaded and 6 virus-infected) grown from two patients (TI006 and TI365) were obtained from the GEO database (accession number GSE117911) [11]. GEO database was also used to obtain the FASTQ files for the rotavirus-infected jejunal-HIEs study (accession number GSE90796). A total of 4 HIEs samples (2 mock and 2 treated) obtained from HIEs grown from two patients (J2 and J11) were deposited under this accession number [12]. Finally, for the astrovirus-infected duodenal-HIEs study FASTQ files were obtained from a total of 6 HIEs samples (3 mock-treated and 3 virus-infected) grown from one patient (D124) [10]. The metadata associated with each one of the samples used in this study is listed in **Supplemental Table 1**.

### Differential expression analysis

Transcript abundances from pseudoalignments (provided in transcripts-per-million) were generated for each RNA-seq dataset using Kallisto [26] and used to generate matrices with estimated gene counts for each one of the RNA-seq datasets re-analyzed. It was taken into account whether the publicly available RNA-seq data was generated from paired-end or single-end Illumina sequencing.

Estimated gene count matrices were then generated for each RNA-seq dataset from transcript abundances using the package tximport [27]. Subsequently we independently filtered out weakly expressed genes by calculating a similarity index among biological replicates using the HTSFilter method which seeks to identify whether genes tend to either have normalized counts less than or equal to the cutoff value in all samples (filtered genes) or greater than the cutoff value in all samples (non-filtered genes) [28]. The set of genes used for differential expression analysis for each RNA-seq dataset consisted then of non-weakly expressed genes that had an annotation in the Entrez Molecular Sequence Database System [29] and a symbol in HUGO Gene Nomenclature Committee (HGNC) [30].

Filtered estimated gene count matrices generated for each RNA-seq dataset were normalized (median of ratios) with the package DESeq2 [31] and subjected to differential expression analysis using the alternative shrinkage estimator ashr [17] to control for false discovery rates (FDR), and effect sizes. A cutoff of an actual fold-change of at least 1.5 (log_2_-FC > 0.58) with a false discovery rate (FDR) cutoff of 1% (adjusted p-value < 0.01) was used to determine whether a gene was differentially expressed. Volcano plots were generated to display the list of DEG for each RNA-seq dataset using the R package EnhancedVolcano [32].

Estimated gene count matrices were also used for exploratory analysis of the dimensionality of the data [33]. Dimensionality reduction was done via two techniques: Principal Component Analysis (PCA) and T-Distributed Stochastic Neighbouring Entities (t-SNE). Each RNA-seq dataset was explored individually to determine the variance within each dataset due to tissue origin of HIEs and treatment (mock-treatment versus viral infection). PCA plots and t-SNE plots were generated using the packages factoextra and Rtsne.

### Over-representation analysis

Over-representation analysis was used to determine whether known biological processes (BP) were enriched in a list of differentially expressed genes (DEGs) for each one of the RNA-seq datasets analyzed. The package clusterprofiler [34] was used for over-representation analysis with strict q-value cutoff of < 0.001 to control for false discovery rate (FDR) using the Benjamini–Hochberg (BH) step-up procedure. Visualization of DEG assigned to each BP category was done using the package ComplexHeatmap [35].

## Code Availability

All the scripts used to generate the figures in this study are available at: https://gitlab.com/ricieza/HIEs-GI-viruses-comparative-transcriptomic.git. The datasets that have been used in this study are publicly available from the GEO and EMBL-EBI indicated above.

---

**Supplemental Figure 1.**
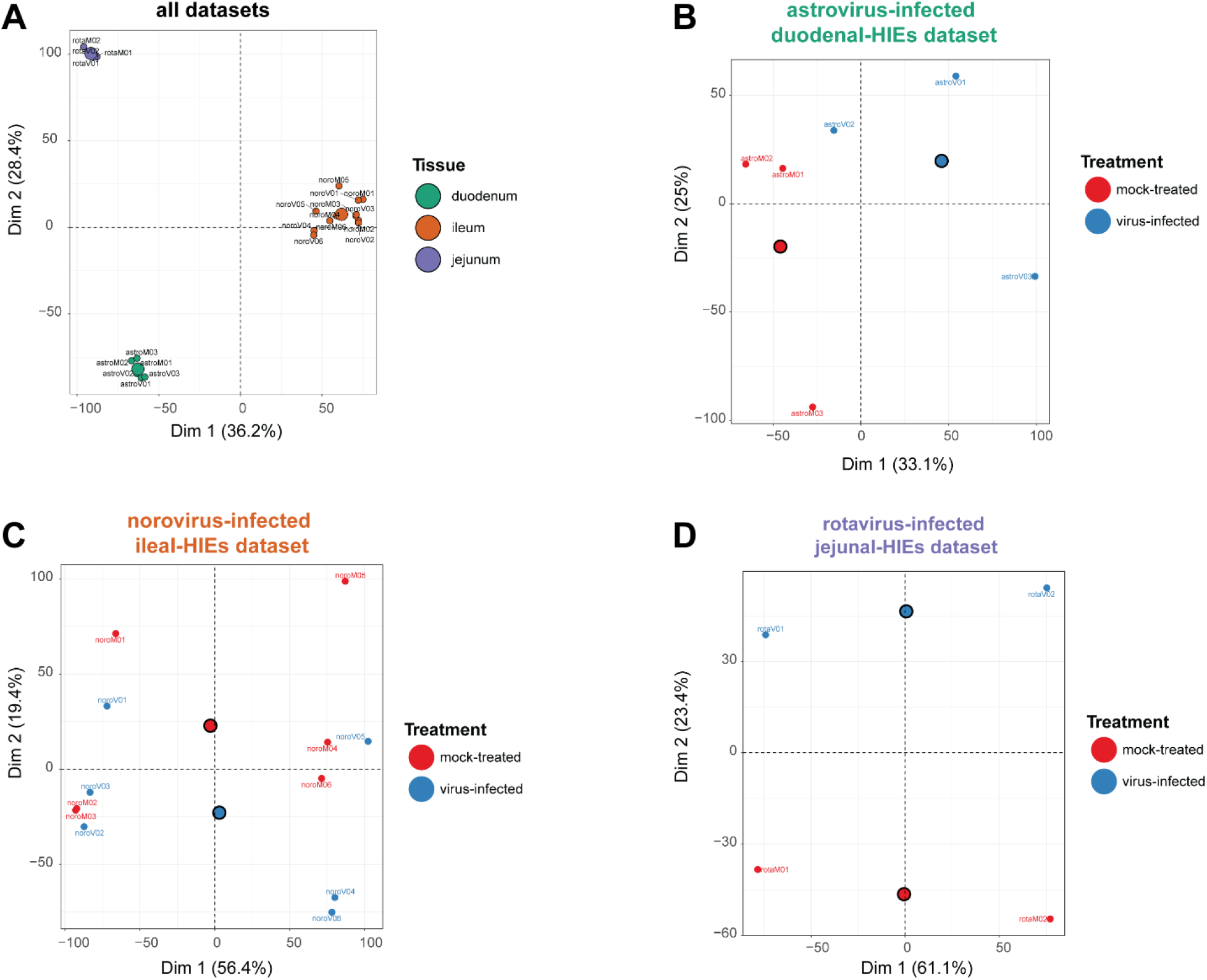
Samples across the three datasets showed divergence due to tissue origin of the HIE line and due to viral infection when looking at each dataset individually. Nonlinear dimensionality reduction with Principal Component Analysis (PCA) of the 16 samples belonging to the three datasets analyzed. Samples are colored by the segment of the small intestine that was used to grow HIE and each one of the datasets analyzed correspond to a different small intestinal segment (duodenum, ileum and jejunum). The PCA plot shows that clustering is primarily driven by the tissue origin of HIE **(A)**. Nonlinear dimensionality reduction with PCA for each individual dataset (6, 12 and 4 samples for astrovirus-infected **(B)**, norovirus-infected **(C)** and rotavirus-infected **(D)** HIEs datasets respectively) shows clustering of the samples due to treatment (mock-treated versus virus-infected) as well as due to HIE line since in the norovirus-infected and rotavirus-infected HIE datasets more than one HIE line were used. PCA plots were generated with the factoextra package in R.

## Supplemental Tables

**Supplemental Table 1: Sequence read archive (SRA) list of all samples analyzed in this study**. FASTQ files from three publicly available RNA-seq datasets were generated from SRA files for downstream differential expression analysis.

**Supplemental Table 2: Non-weakly expressed genes shared in all RNA-seq datasets and partially shared across datasets**. List of non-weakly expressed genes identified in astrovirus-infected, norovirus-infected and rotavirus-infected HIE.

**Supplemental Table 3: Uniquely differentially expressed genes (DEGs) for each RNA-seq dataset analyzed**. List of DEGs identified in a single RNA-seq dataset re-analyzed.

## References

1. Oude Munnink BB, van der Hoek L: Viruses Causing Gastroenteritis: The Known, The New and Those Beyond. Viruses 2016, 8(2).

2. Hall AJ, Rosenthal M, Gregoricus N, Greene SA, Ferguson J, Henao OL, Vinje J, Lopman BA, Parashar UD, Widdowson MA: Incidence of acute gastroenteritis and role of norovirus, Georgia, USA, 2004-2005. Emerg Infect Dis 2011, 17(8):1381–1388.

3. Ahmed SM, Hall AJ, Robinson AE, Verhoef L, Premkumar P, Parashar UD, Koopmans M, Lopman BA: Global prevalence of norovirus in cases of gastroenteritis: a systematic review and metaanalysis. Lancet Infect Dis 2014, 14(8):725–730.

4. Crawford SE, Ramani S, Tate JE, Parashar UD, Svensson L, Hagbom M, Franco MA, Greenberg HB, O’Ryan M, Kang G et al: Rotavirus infection. Nat Rev Dis Primers 2017, 3:17083.

5. Vu DL, Bosch A, Pinto RM, Guix S: Epidemiology of Classic and Novel Human Astrovirus: Gastroenteritis and Beyond. Viruses 2017, 9(2).

6. Czerwinski M HE, Yu-Hwai Tsai, Angeline Wu, Qianhui Yu, Josh Wu, Katherine D. Walton, Caden Sweet, Charlie Childs, Ian Glass, Barbara Treutlein, J. Gray Camp, Spence JR: In vitro and in vivo development of the human intestinal niche at single cell resolution. 2020.

7. Lauder E KK, Schmidt TM, Golob JL: Organoid-derived adult human colonic epithelium responds to co-culture with a probiotic strain of Bifidobacterium longum. Systems Biology 2020.

8. Blutt SE, Crawford SE, Ramani S, Zou WY, Estes MK: Engineered Human Gastrointestinal Cultures to Study the Microbiome and Infectious Diseases. Cell Mol Gastroenterol Hepatol 2018, 5(3):241–251.

9. Kolawole AO, Wobus CE: Gastrointestinal organoid technology advances studies of enteric virus biology. PLoS Pathog 2020, 16(1):e1008212.

10. Kolawole AO, Mirabelli C, Hill DR, Svoboda SA, Janowski AB, Passalacqua KD, Rodriguez BN, Dame MK, Freiden P, Berger RP et al: Astrovirus replication in human intestinal enteroids reveals multi-cellular tropism and an intricate host innate immune landscape. PLoS Pathog 2019, 15(10):e1008057.

11. Hosmillo M, Chaudhry Y, Nayak K, Sorgeloos F, Koo BK, Merenda A, Lillestol R, Drumright L, Zilbauer M, Goodfellow I: Norovirus Replication in Human Intestinal Epithelial Cells Is Restricted by the Interferon-Induced JAK/STAT Signaling Pathway and RNA Polymerase II-Mediated Transcriptional Responses. mBio 2020, 11(2).

12. Saxena K, Simon LM, Zeng XL, Blutt SE, Crawford SE, Sastri NP, Karandikar UC, Ajami NJ, Zachos NC, Kovbasnjuk O et al: A paradox of transcriptional and functional innate interferon responses of human intestinal enteroids to enteric virus infection. Proc Natl Acad Sci U S A 2017, 114(4):E570–E579.

13. Sudmant PH, Alexis MS, Burge CB: Meta-analysis of RNA-seq expression data across species, tissues and studies. Genome Biol 2015, 16:287.

14. Fasterius E, Al-Khalili Szigyarto C: Analysis of public RNA-sequencing data reveals biological consequences of genetic heterogeneity in cell line populations. Sci Rep 2018, 8(1):11226.

15. Lin SC, Qu L, Ettayebi K, Crawford SE, Blutt SE, Robertson MJ, Zeng XL, Tenge VR, Ayyar BV, Karandikar UC et al: Human norovirus exhibits strain-specific sensitivity to host interferon pathways in human intestinal enteroids. Proc Natl Acad Sci U S A 2020, 117(38):23782–23793.

16. Baccarella A, Williams CR, Parrish JZ, Kim CC: Empirical assessment of the impact of sample number and read depth on RNA-Seq analysis workflow performance. BMC Bioinformatics 2018, 19(1):423.

17. Stephens M: False discovery rates: a new deal. Biostatistics 2017, 18(2):275–294.

18. He L, Wang B, Li Y, Zhu L, Li P, Zou F, Bin L: The Solute Carrier Transporter SLC15A3 Participates in Antiviral Innate Immune Responses against Herpes Simplex Virus-1. J Immunol Res 2018, 2018:5214187.

19. McNab F, Mayer-Barber K, Sher A, Wack A, O’Garra A: Type I interferons in infectious disease. Nat Rev Immunol 2015, 15(2):87–103.

20. Martin SK, Tomida J, Wood RD: Disruption of DNA polymerase zeta engages an innate immune response. Cell Rep 2021, 34(8):108775.

21. Pervolaraki K, Stanifer ML, Munchau S, Renn LA, Albrecht D, Kurzhals S, Senis E, Grimm D, Schroder-Braunstein J, Rabin RL et al: Type I and Type III Interferons Display Different Dependency on Mitogen-Activated Protein Kinases to Mount an Antiviral State in the Human Gut. Front Immunol 2017, 8:459.

22. Su R, Shereen MA, Zeng X, Liang Y, Li W, Ruan Z, Li Y, Liu W, Liu Y, Wu K et al: The TLR3/IRF1/Type III IFN Axis Facilitates Antiviral Responses against Enterovirus Infections in the Intestine. mBio 2020, 11(6).

23. Ioannidis I, Ye F, McNally B, Willette M, Flano E: Toll-like receptor expression and induction of type I and type III interferons in primary airway epithelial cells. J Virol 2013, 87(6):3261–3270.

24. McCartney SA, Thackray LB, Gitlin L, Gilfillan S, Virgin HW, Colonna M: MDA-5 recognition of a murine norovirus. PLoS Pathog 2008, 4(7):e1000108.

25. Perng YC, Lenschow DJ: ISG15 in antiviral immunity and beyond. Nat Rev Microbiol 2018, 16(7):423–439.

26. Bray NL, Pimentel H, Melsted P, Pachter L: Near-optimal probabilistic RNA-seq quantification. Nat Biotechnol 2016, 34(5):525–527.

27. Soneson C, Love MI, Robinson MD: Differential analyses for RNA-seq: transcript-level estimates improve gene-level inferences. F1000Res 2015, 4:1521.

28. Rau A, Gallopin M, Celeux G, Jaffrezic F: Data-based filtering for replicated high-throughput transcriptome sequencing experiments. Bioinformatics 2013, 29(17):2146–2152.

29. Gibney G, Baxevanis AD: Searching NCBI databases using Entrez. Curr Protoc Bioinformatics 2011, Chapter 1:Unit 13.

30. Naming human genes. Nat Genet 2020, 52(8):751.

31. Love MI, Huber W, Anders S: Moderated estimation of fold change and dispersion for RNA-seq data with DESeq2. Genome Biol 2014, 15(12):550.

32. Blighe K RS, Lewis M: EnhancedVolcano: Publication-ready volcano plots with enhanced colouring and labeling. 2018.

33. Simmons S, Peng J, Bienkowska J, Berger B: Discovering What Dimensionality Reduction Really Tells Us About RNA-Seq Data. J Comput Biol 2015, 22(8):715–728.

34. Yu G, Wang LG, Han Y, He QY: clusterprofiler: an R package for comparing biological themes among gene clusters. OMICS 2012, 16(5):284–287.

35. Gu Z, Eils R, Schlesner M: Complex heatmaps reveal patterns and correlations in multidimensional genomic data. Bioinformatics 2016, 32(18):2847–2849.

